# The cost of being stable: Trade-offs between effort and stability across a landscape of redundant motor solutions

**DOI:** 10.1101/477083

**Authors:** M. Hongchul Sohn, Lena H. Ting

**Author notes:** Current Address: Department of Biomedical Engineering, Northwestern University, Evanston, Illinois, United States of America (M. Hongchul Sohn). Current Address: The Shirley Ryan AbilityLab, Chicago, Illinois, United States of America (M. Hongchul Sohn). Corresponding author (LHT).

## Abstract

Current musculoskeletal modeling approaches cannot account for variability in muscle activation patterns seen across individuals, who may differ in motor experience, motor training, or neurological health. While musculoskeletal simulations typically select muscle activation patterns that minimize muscular effort, and generate unstable limb dynamics, a few studies have shown that maximum-effort solutions can improve limb stability. Although humans and animals likely adopt solutions between these two extremes, we lack principled methods to explore how effort and stability shape how muscle activation patterns differ across individuals. Here we characterized trade-offs between muscular effort and limb stability in selecting muscle activation patterns for an isometric force generation task in a musculoskeletal model of the cat hindlimb. We define effort as the sum of squared activation across all muscles, and limb stability by the maximum real part of the eigenvalues of the linearized musculoskeletal system dynamics, with more negative values being more stable. Surprisingly, stability increased rapidly with only small increases in effort from the minimum-effort solution, suggesting that very small amounts of muscle coactivation are beneficial for postural stability. Further, effort beyond 40% of the maximum possible effort did not confer further increases in stability. We also found multiple muscle activation patterns with equivalent effort and stability, which could underlie variability observed across individuals with similar motor ability. Trade-off between muscle effort and limb stability could underlie diversity in muscle activation patterns observed across individuals, disease, learning, and rehabilitation.

**Author summary:** Current computational musculoskeletal models select muscle activation patterns that minimize the amount of muscle activity used to generate a movement, creating unstable limb dynamics. However, experimentally, muscle activation patterns with various level of co-activation are observed for performing the same task both within and across individuals that likely help to stabilize the limb. Here we show that a trade-off between muscular effort and limb stability across the wide range of possible muscle activation patterns for a motor task could explain the diversity of muscle activation patterns seen across individuals, disease, learning and rehabilitation. Increased muscle activity is necessary to stabilize the limb, but could also limit the ability to learn new muscle activation pattern, potentially providing a mechanism to explain individual-specific muscle coordination patterns in health and disease. Finally, we provide a straightforward method for improving the physiological relevance of muscle activation pattern and musculoskeletal stability in simulations.

## Introduction

Little is known about the landscape of redundant motor solutions for sensorimotor tasks, hindering our ability to interpret variability in muscle activation patterns seen across individuals, who may differ in experience, training, or neurological health. In general, we refer to motor solutions as the spatiotemporal pattern of motor neuron activity that are reflected in electromyographic (EMG) signals during posture and movement. More specifically, we have demonstrated that individual-specific motor modules–defined as library of fixed spatial patterns of muscle activity that execute a motor task–can vary considerably across individuals. Experimental EMG recordings reveal variations across individuals in the degree to which muscles are co-activated in motor modules that perform similar tasks [1, 2]. Further, differences in motor module structure are amplified when examining individual across the spectrum of motor ability: long-term motor training, such as in ballet dancers, tends to decrease the number of muscles used to execute a given task [3], whereas neurological impairments tend to increase the co-activation of muscles used to perform gait and balance tasks [4-8]. For reactive balance tasks in cats, we explicitly showed individual-specific differences in motor modules that receive similar neural commands and produce equivalent motor outputs in healthy, trained, animals [9]. Through analysis of musculoskeletal models, we found the biomechanics of the limb and task requirements to impose few constraints on feasible muscle activation patterns [10-13]. These results indicate that there is ample room in the nullspace, i.e. set of possible motor solutions, from which individuals may employ motor modules that deviate from the muscle coordination pattern predicted by single optimality criteria such as minimizing muscle effort [14-16]. However, we understand little about what might drive the selection of particular individual-specific patterns within the nullspace of a given task.

While minimizing muscular effort has been established as a principle driving muscle activation patterns for movement, increasing muscle effort to stabilize the limb is also an important principle of motor control that may be especially pertinent in understanding impaired motor control in neurological disorders. Muscular effort is often used as a cost to be minimized in motor tasks, including reaching and walking [17-19]. However, the level of muscle co-contraction can differ across instances of the same motor task, increasing both limb stability and muscular effort [20-25]. Such open-loop stability conferred by active muscles is required to slow the intrinsic dynamics of the limb because neural control of movement via sensory feedback is subject to long conduction delays [26-28]. Specifically in reactive balance control, muscle force responses elicited by perturbation are delayed by ~70 and ~150 ms in cats and humans, respectively [29]. Background muscle activation in quiet standing thus provides critical intrinsic stability sufficient to reduce postural changes during the response latency period [28, 30]. Experimentally, individuals can modify motor solutions to modulate limb or whole-body stability on a trial-by-trial basis [31] as well as over long-term motor training [3]. Elevated muscle co-contraction common in individuals with impaired neural control of movement [32] likely increases limb and body stability as a compensatory strategy for impaired neural feedback control. Although limb and body stability is an important functional property for motor control, how stability consideration affects muscular effort in the selection of muscle activation patterns is not understood.

Due to the lack of computational methods to characterize the functional landscape of redundant motor solutions, explicit trade-offs between muscular effort and limb stability in motor solutions have not been examined in musculoskeletal models. Minimum-effort criteria [33-35] often yield unstable musculoskeletal dynamics where a small disturbance or numerical error lead to unrecoverable deviation from the desired state trajectory in forward simulations [36-38]. On the other hand, maximum-effort solutions can substantially improve stability, as measured by endpoint stiffness of human arm estimated during postural task [39]. In general, however, the relationship between muscular effort and stability along the continuum between the two extrema– where actual motor solutions most likely lie–is unknown. By applying Lyapunov stability theory to a musculoskeletal model of that cat hindlimb, we demonstrated that limb stability can be modulated by the selection of diverse patterns of muscle activation pattern within the nullspace of equivalent solutions in terms of force output [40]. Stability was evaluated by computing the eigenvalues of the linearized system dynamics, incorporating both rigid body dynamics and force-length and force-velocity properties of activated muscles [36, 40]. Our prior work only examined stability of randomly sampled solutions throughout the nullspace, without consideration for muscular effort. To our knowledge, no study has characterized the functional landscape of stability along the trajectory between minimum- and maximum- effort solutions where effort level is systematically varied. Here our goal was to quantitatively express trade-offs between effort and stability to understand how motor solution might differ across tasks and individuals.

Given that individuals may seek to modulate muscular effort and/or limb stability [41, 42], it is also not known whether one could learn a better motor solution by making small changes to a muscle activation pattern within the nullspace of motor solutions. The observation that individuals maintain characteristic motor solutions, e.g. habitual [43-45] or pathological [46-48], suggests that it may be difficult to find nearby solutions that improve the functional properties while also satisfying task requirements. In particular, without a priori knowledge of what direction of change in muscle space would satisfy the task constraint, how easy is it to find a neighboring solution that satisfies the tasks constraint and desired changes in the functional properties? Describing the local landscape around particular motor solutions with respect to effort and stability may help explaining individual differences in the ability to learn a new motor solution. For example, particular motor solution may have limited access to neighboring solutions with better functional properties depending on where they lie initially in the functional property space, as well as the extent to which they explore the null space. Thus, it still remains unknown, given a particular solution, how much of muscular effort is required to meaningfully increase limb stability, or how much effort could be reduced without sacrificing limb stability. To address this issue, we explored the local landscape around different motor solutions in terms of the changes in their functional properties.

It is also possible that motor solutions differ across individuals because the nullspace of motor solutions has many local minima in terms of their functional properties. While previous works have identified wide bounds in nullspace of motor solutions [10, 12, 13], and spatial structure in multitude of muscle activation patterns that lie within that space [49], it remains unknown whether such redundancy also generate functional equivalence in many different solutions. The existence of functionally equivalent motor solutions could explain how two people could arrive on different solutions yet with seemingly similar performance [2, 9, 50, 51]. Previous work has shown that individuals could learn two non-adjacent motor solutions to a task and have difficulty finding the other solution unless it is explicitly demonstrated [44, 45, 52]. It is not clear how many such motor solutions may provide “good enough” solutions exist for a given task [53], and whether they can differ substantially in the muscle activation patterns. Therefore, we also tested whether similar functional properties could be achieved with widely diverging muscle activation pattern.

In summary, our overall objective was to explore the nullspace of motor solutions for a quasi-static force production task (Fig 1A) with respect to the two functional properties of effort and stability using a 3D musculoskeletal model of the cat hindlimb (Fig 1B). Specifically, we examined 1) whether functional landscape of stability along the trajectory between minimum- and maximum- effort solutions indeed express explicit trade-off between effort and stability; 2) whether local search around viable muscle activation patterns is a feasible method of searching for motor solutions with better functional properties and 3) whether qualitatively different muscle activation patterns have near-equivalent functional properties, consistent with the existence of multiple local minima within the nullspace of motor solutions. We found that stability increased rapidly with only small increases in effort from the minimum-effort solution, suggesting that very small amounts of muscle co-activation are beneficial for postural stability. Further, effort beyond 40% of the maximum possible effort did not confer further increases in stability. We also found multiple muscle activation patterns with equivalent effort and stability, which could underlie variability observed across individuals with similar motor ability. Our work provides evidence that trade-off between muscle effort and limb stability could underlie diversity in muscle activation patterns observed across individuals, disease, learning, and rehabilitation.

**Fig 1.**
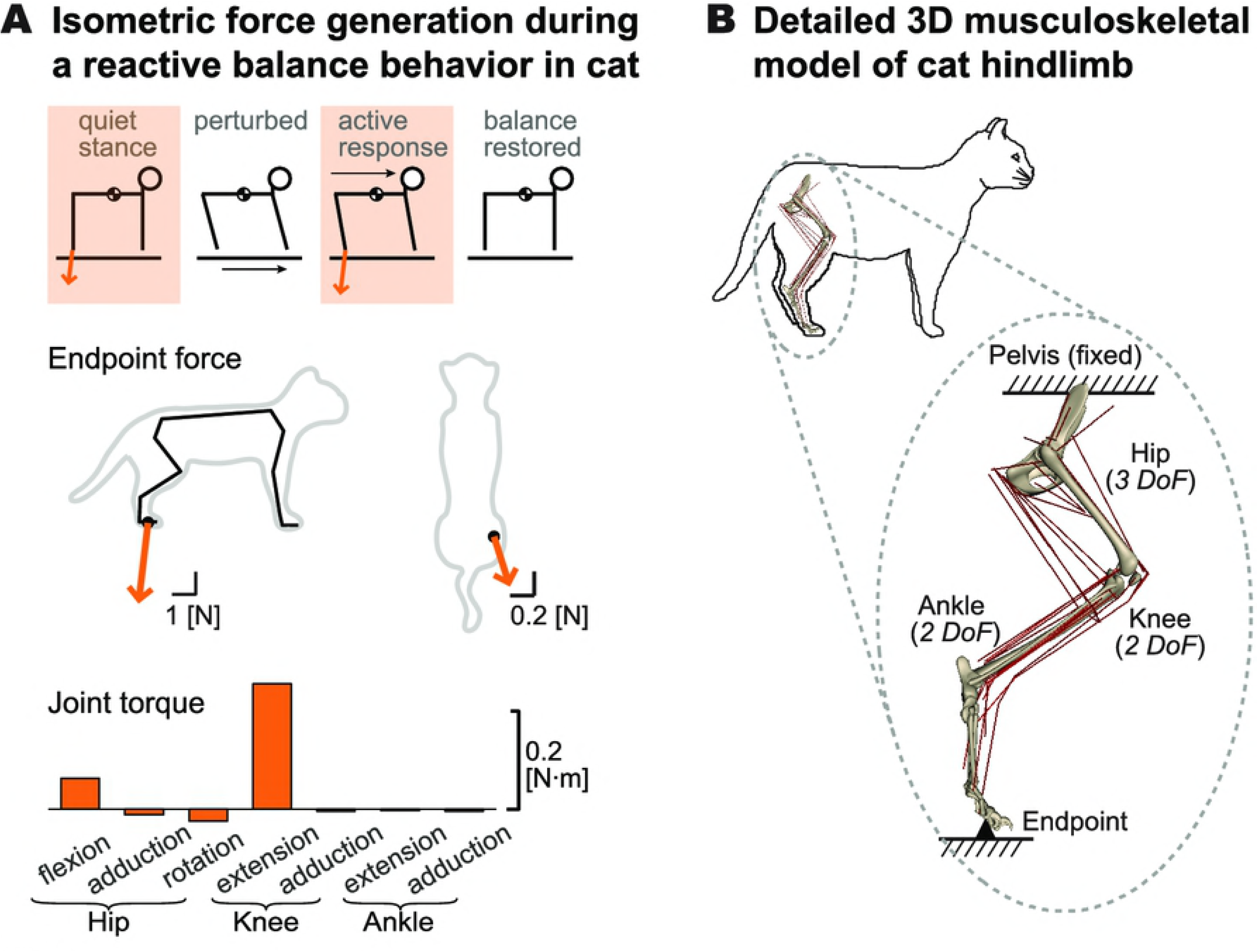
Experimentally-derived motor task and musculoskeletal model. (A) An extensor force vector typically observed during quiet standing and active postural responses to perturbation [9] (top) was used as the target endpoint force vector (middle). The target endpoint force primarily required producing knee extension torque and hip flexion torque (bottom). (B) The model included seven degrees of freedom at the anatomical joints (3 at the hip, 2 at the knee, and 2 at the ankle) and 31 Hill-type muscle actuators (see Table 1 for list and abbreviations). The pelvis was fixed to the ground and the endpoint, defined at the metatarsophalangeal joint, was connected to the ground via gimbal joint.

## Results

### Trade-off between effort and stability along a continuum of solutions in the nullspace

We found dramatic differences in the effort and stability of the minimum- and maximum-effort solutions for generating an extensor force typically observed during quiet standing as well as during postural responses to perturbation in cats (Fig 1A). When normalized as percent of the maximum possible effort level (i.e. 100%), the same task could be achieved with the minimum-effort solution at only 8.80% effort of the maximum-effort solution. We then evaluated the stability of each solution using a stability metric defined as the maximum real part of the eigenvalue of the musculoskeletal system (***S***=max{Re(λ)}) when considering the contributions of active muscle to the overall system dynamics; systems are unstable when ***S***>0. As expected [36, 37, 39], the minimum-effort solution (Fig 2A, ‘x’ in light blue) was unstable (***S***=18.9), indicating that the limb configuration would catastrophically collapse in tens of milliseconds (doubling time=36.7ms) when perturbed. On the other hand, the maximum-effort solution (Fig 2A ‘x’ in black) was stable (***S***=-2.25), allowing it to maintain the postural configuration and reject external perturbations, consistent with prior studies [20, 22].

**Table 1.** Muscles included in the hindlimb model and abbreviations.

| Name | Abbreviation | Name | Abbreviation |
| --- | --- | --- | --- |
| *Adductor femoris* | adf | *Plantaris* | plan |
| *Adductor longus* | adl | *Iliopsoas* | psoas |
| *Biceps femoris anterior* | bfa | *Peroneus tertius* | pt |
| *Biceps femoris posterior* | bfp | *Pyriformis* | pyr |
| *Extensor digitorum longus* | edl | *Quadratus femoris* | qf |
| *Flexor digitorum longus* | fdl | *Rectus femoris* | rf |
| *Flexor hallicus longus* | fhl | *Sartorius* | sart |
| *Gluteus maximus* | gmax | *Semimembranosus* | sm |
| *Gluteus medius* | gmed | *Soleus* | sol |
| *Gluteus minimus* | gmin | *Semitendinosus* | st |
| *Gracilis* | grac | *Tibialis anterior* | ta |
| *Lateral gastrocnemius* | lg | *Tibialis posterior* | tp |
| *Medial gastrocnemius* | mg | *Vastus intermedius* | vi |
| *Peroneus brevis* | pb | *Vastus lateralis* | vl |
| *Pectineus* | pec | *Vastus medialis* | vm |
| *Peroneus longus* | pl |  |  |

157 Muscles are listed in alphabetic order.

**Fig 2.**
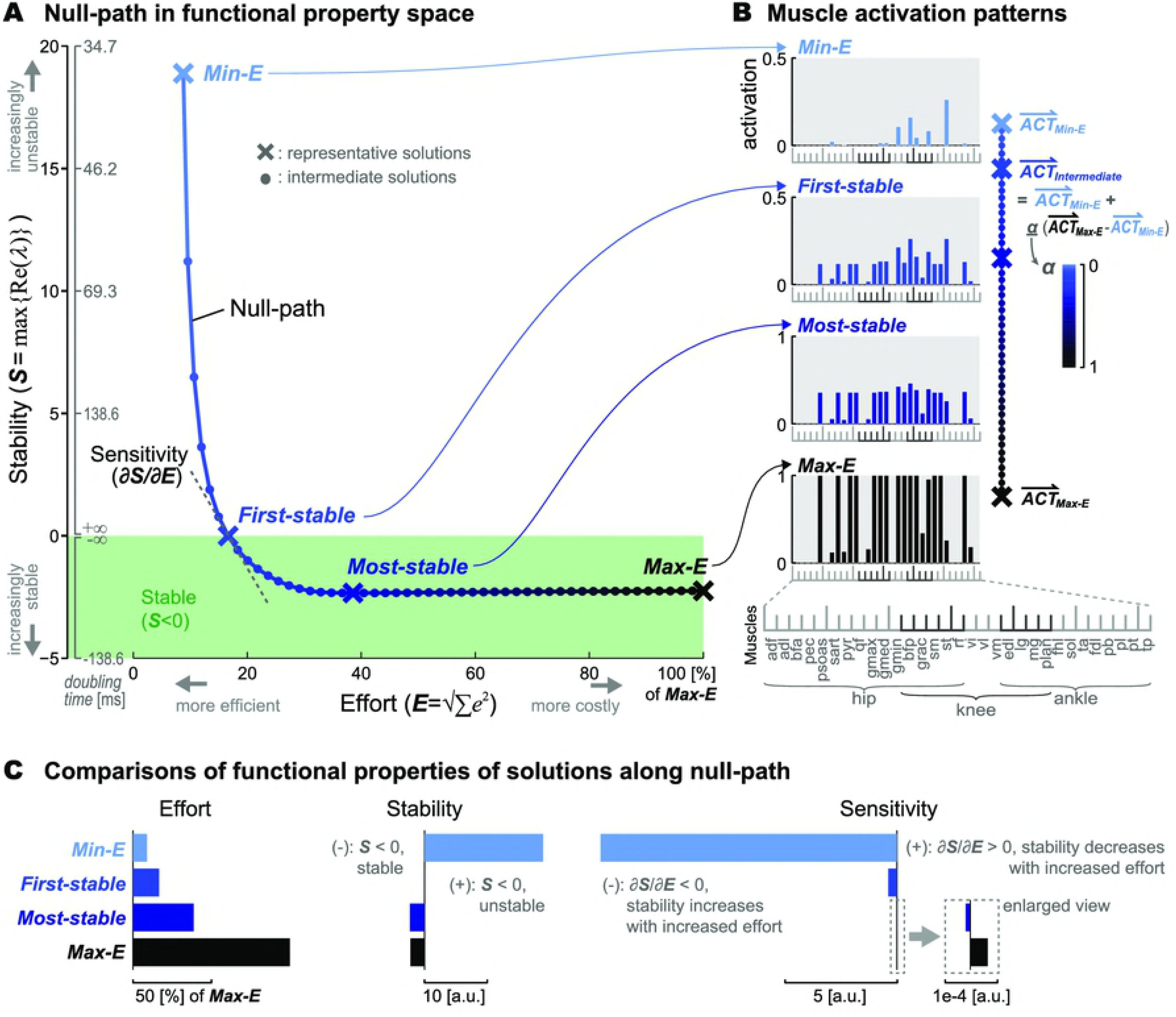
Effort and stability along a continuum of solutions in the null space. (A) Muscle activation patterns with same task output and different effort and stability were evaluated in the functional property space. The minimum-effort solution (Min-E, ‘x’ in light blue) was unstable, while the maximum-effort solution (Max-E, ‘x’ in black) was stable. Stability rapidly improved with increased effort in the low-effort region, e.g. from Min-E to the first solution (First-stable, ‘x’ in blue) on the null-path that became stable (***S***<0). Stability did not further improve beyond moderate level of effort (~40%), e.g. the most stable solution on the null-path (Most-stable, ‘x’ in dark blue). (B) Muscle activation patterns along the nullspace between the minimum- (Min-E, ‘x’ in light blue) and maximum effort solutions (Max-E, ‘x’ in black) constituting the null-path. Intermediate solutions (e.g. middle two rows) were identified by linearly combining the minimum- and maximum-effort solutions, where the difference between the two solutions were linearly scaled (α, values from 0 to 1 represented with color gradient from light blue to black) through the null muscle activation space. (C) Effort (left), stability (middle), and sensitivity of stability to change in effort (right) for the four representative solutions: minimum-effort (light blue), first-stable (blue), most-stable (dark blue), and maximum-effort (black) solutions. Sensitivity was defined as the instantaneous slope of the null-path at each solution, calculated by numerical differentiation. [a.u.] stands for arbitrary unit.

We found a reciprocal relationship between effort and stability when stability was examined along the continuum of solutions between the two the extremes identified by the minimum- and maximum-effort solutions (Fig 2A). We identified the set of muscle activation patterns along a null-path, defined as the linear combinations of minimum-effort and maximum-effort solutions (Fig 2B) for producing the same endpoint force vector. As such, the minimum-effort solution was the most unstable (greatest ***S***) solution among the 51 solutions on this null-path. As effort increased, the stability conferred by the muscle activation improved (***S*** decreased).

To further characterize solutions along the null-path, we computed the effort and stability of the solution where the path transitioned from unstable to stable solutions (i.e. from ***S***>0 to ***S***<0), and the point of maximum stability along the path. The first solution on the null-path that became stable (Fig 2A, ‘x’ in blue) had only 8% increase in effort from the minimum-effort solution (**S**=-0.0066 and ***E***=16.6%). The most stable solution on the null-path (Fig 2A, ‘x’ in dark blue, ***S***=-2.34) required only a moderate level of effort, ***E***=38.6% compared to the maximum-effort solution.

The slope of the null-path indicates the sensitivity of the system’s postural stability to changes in muscular effort, which is very steep for low effort solutions and insensitive for solution of greater than ~40% effort (Fig 2A). The most sensitive region was between the minimum-effort solution and the first stable solution, where small increase in effort resulted in rapid improvement in stability (e.g. Min-E in Fig 2C). From the most stable solution to the maximum-effort solution, however, stability was almost insensitive to change in effort (e.g. Most-stable in Fig 2C). More importantly, beyond the most stable solution (***E***=38.6%), further increases in effort decreased stability (e.g. Max-E in Fig 2C).

### Local landscape of effort and stability vary along the null-path

Because the null-path only identifies solutions along one particular direction in the high dimensional nullspace (24, for our musculoskeletal model), we further explored the local landscape around 51 solutions along the null-path (Fig 3). We were particularly interested in whether applying local perturbations to the pattern of muscle activation of a particular solution on the null-path create a set of solutions that preserve the characteristics of the null path in the functional property space. To this end, each solution on the null-path was used to as a seed solution (Fig 3A), from which we found multiple set of neighboring solutions randomly perturbed in muscle space. To determine the size of perturbation, we first identified feasible range of activation for each muscle [13] (Fig 3A, gray boxes). We then generated three sets of random perturbations that were of 5, 10, and 25% of the feasible range that were normally distributed about the activation of each muscle in the seed solution (Fig 3B, perturbation distribution). To ensure that these patterns satisfied the force generation task, we projected the perturbed muscle activation patterns to the linear solution manifold (Fig 3B, solution distribution).

**Fig 3.**
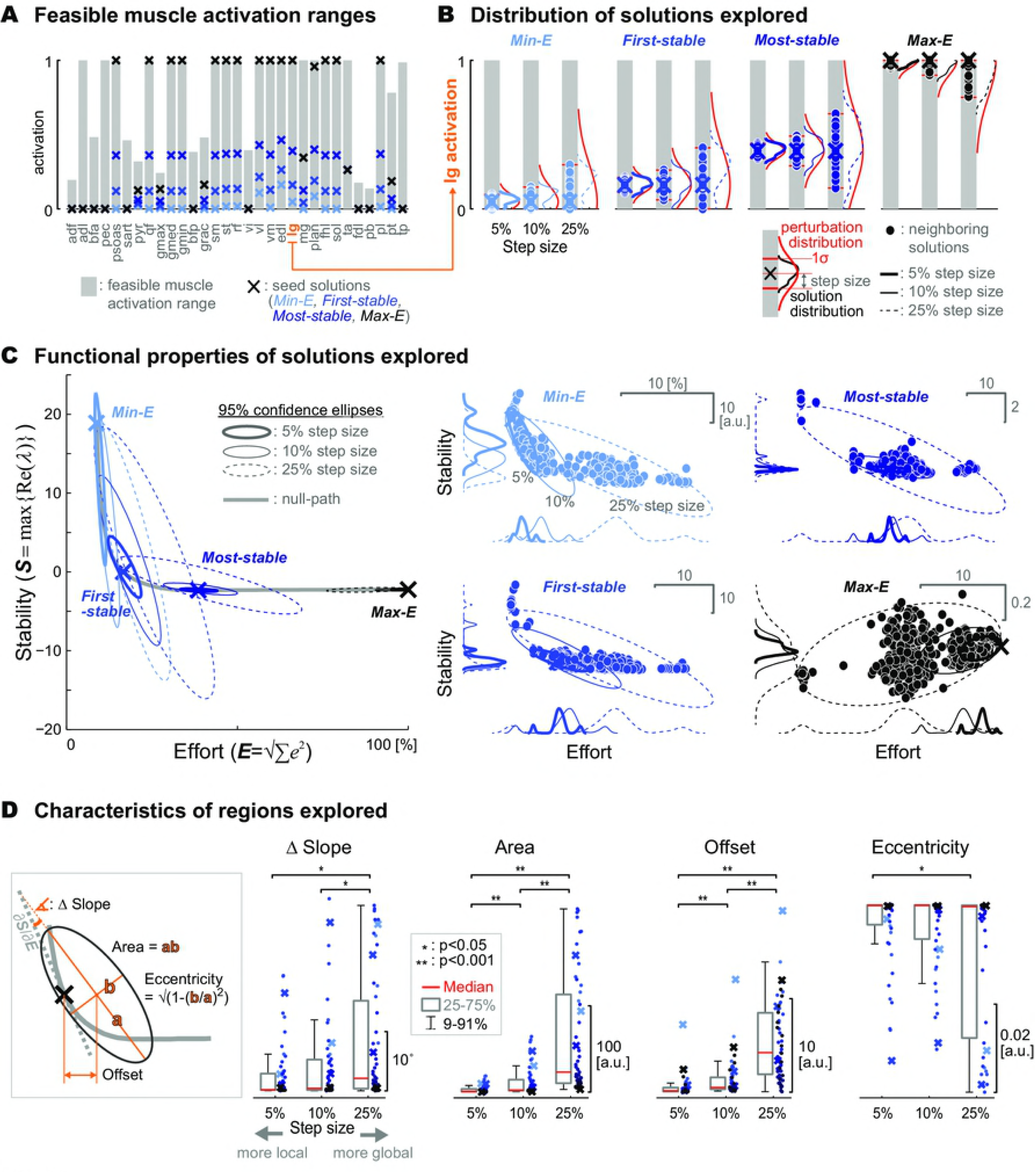
Local landscape of effort and stability around the solutions on the null-path. (A) Feasible muscle activation ranges (gray bar) for satisfying the force generation task were identified for each muscle [13]. Each of the 51 solutions on the null-path (four examples in ‘x’) were used as a seed, from which a perturbation of different size relative to the feasible muscle activation range was applied. (B) Examples of neighboring solutions explored: activation of lateral gastrocnemius (lg) for the four representative solutions. Three sets of random perturbations, normally distributed (red) about the activation of the seed solution (‘x) with step size (=1σ) corresponding to 5, 10, and 25% of the feasible range were generated. Neighboring solutions (dots) were then found by projecting theses perturbed patterns to the solution manifold. Resulting distributions are shown in thick solid line for 5% step size, thin solid line for 10% step size, and dotted line for 25% steps size. (C) Two-dimensional 95% confidence ellipses were found for each set of neighboring solutions, i.e., for each seed solution (four representative examples in ‘x’) and steps size (5%, 10%, and 25% in thick solid, thin solid, and dotted line, respectively). The ellipse was centered at the mean effort and stability, with major axes in the direction of maximum covariance of neighboring solutions in the functional property space. The local landscape explored by the neighboring solutions closely followed the null-path (gray line). Changes in functional property occurred mainly in stability near low effort region, e.g. the minimum-effort (‘x’ in light blue) solution, while it was very difficult to change stability near high effort region, e.g. the maximum-effort solution (‘x’ in black). Actual neighboring solutions are shown on the right panels (note different scales), with distributions shown in thick solid, thin solid, and dotted lines for 5%, 10%, and 25% step size, respectively. [a.u.] stands for arbitrary unit. (D) With increased step size, the orientation of the 95% ellipse deviated more from the slope of the original seed solution on the null path (ΔSlope), and larger region (Area) with more different effort and stability (Offset) were found. Explored solutions also spanned wider range of directions around the seed solution (Eccentricity) with increased step size.

Local searches around seed solutions on the null-path preserved the functional properties of the seed solution, with solutions becoming spread across the functional property space as step size increased. We characterized the set of solutions around each seed point using a two-dimensional ellipse that was centered at the mean, and the axes and radii defined by the covariance of effort and stability values of all corresponding neighboring solutions (Fig 3C). Near the lowest effort region, e.g. the minimum-effort solution (Fig 3C, light blue), it was “easy” to find more stable solutions with small increases in effort. While few solutions that are more unstable (greater ***S***) than the minimum-effort solution could be found (Fig 3C right panel, light blue), most neighboring solutions became more stable (decreased ***S***). However, a larger change in the muscle activation pattern, e.g. 25% step size (Fig 3C, dotted line in light blue), was required to find a stable solution (***S***<0). Local search around the border of marginal stability (***S***=0), e.g. the first-stable solution (Fig 3C, blue), reached both unstable solutions (***S***>0) with lower effort as well as more stable solutions with increased effort, suggesting that even small change in muscle activation pattern may result in destabilization. Beyond the most-stable solution (Fig 3C, dark blue) where the change in stability becomes insensitive to change in effort (i.e., slope near zero), local search also resulted in finding neighboring solutions that mostly varied in effort, without much change in stability. Notably, it was difficult to change stability when searching around the maximum-effort solution (Fig 3C, black).

As step size increased, solutions differed more compared to the seed solution (Fig 3D). Across local searches from all 51 seed solutions, the difference between the major axis of each ellipse and the slope of the null-path for each seed solution increased with step size (ΔSlope; χ^2^(2)=12.4, p=0.002). In particular, the change in orientation of the ellipses from the slope of the null-path were greater with 25% step size, compared to 5% (p=0.002) and 10% (p=0.041). As step size increased, the area (χ^2^(2)=76.1, p=3.05e-17) of the ellipse and the offset of the center from the seed (χ^2^(2)=76.8, p=2.09e-17) also increased, whereas eccentricity decreased (χ^2^(2)=6.85, p=0.033), indicating that solutions that are further from the null-path in the functional property space were explored. The area for the 25% step size search was greater than for the 5% (p=9.56e-10) and 10% (p=7.97e-6) step size, and the area for the 10% step size was greater than that for the 5% step size (p=1.74e-4). Similarly, offset for the 25% step size was greater than for the 5% (p=9.56e-10) and 10% (p=2.30e-6) step size, and area for the 10% step size was greater than that for the 5% step size (p=4.31e-4). Eccentricity of the 95% confidence ellipses for neighboring solutions explored with 25% step size was smaller than the 5% step size (p=0.049).

### Multiple local solutions can have equivalent functional properties

We found multiple solutions that had very similar effort and limb stability near the most-stable solution (Fig 4A, ‘x’ in dark blue) on the null-path, but which differed in their muscle activation patterns. The difference in effort and stability of 166 selected solutions (Fig 4A, gray dots) compared to the most-stable solution were only −0.003±0.2% and −0.02±0.11% (mean±std), respectively. Nevertheless, a wide variety of muscle activation patterns was observed among these 166 nearest solutions. For example, for two solutions that were very close to each other in functional property space (Fig 4A, highlighted in magenta and red), load-sharing [54, 55] between hip extensors (e.g. gmed and gmin, see Table 1 for abbreviations) differed between the two solutions (Fig 4B, top). Hamstring muscles (e.g. sm, st) were highly activated in one solution (Fig 4B, magenta), which resulted in activation of hip flexor/knee extensor (e.g. rf) for torque balance at the knee. However, in the other solution (Fig 4B, red), vm was mainly used for knee extension. Large activation of mg and lg, along with co-activation of sol and ta, were present in one solution (Fig 4B, magenta), whereas the other solution (Fig 4B, red) had no mg and sol activation.

**Fig 4.**
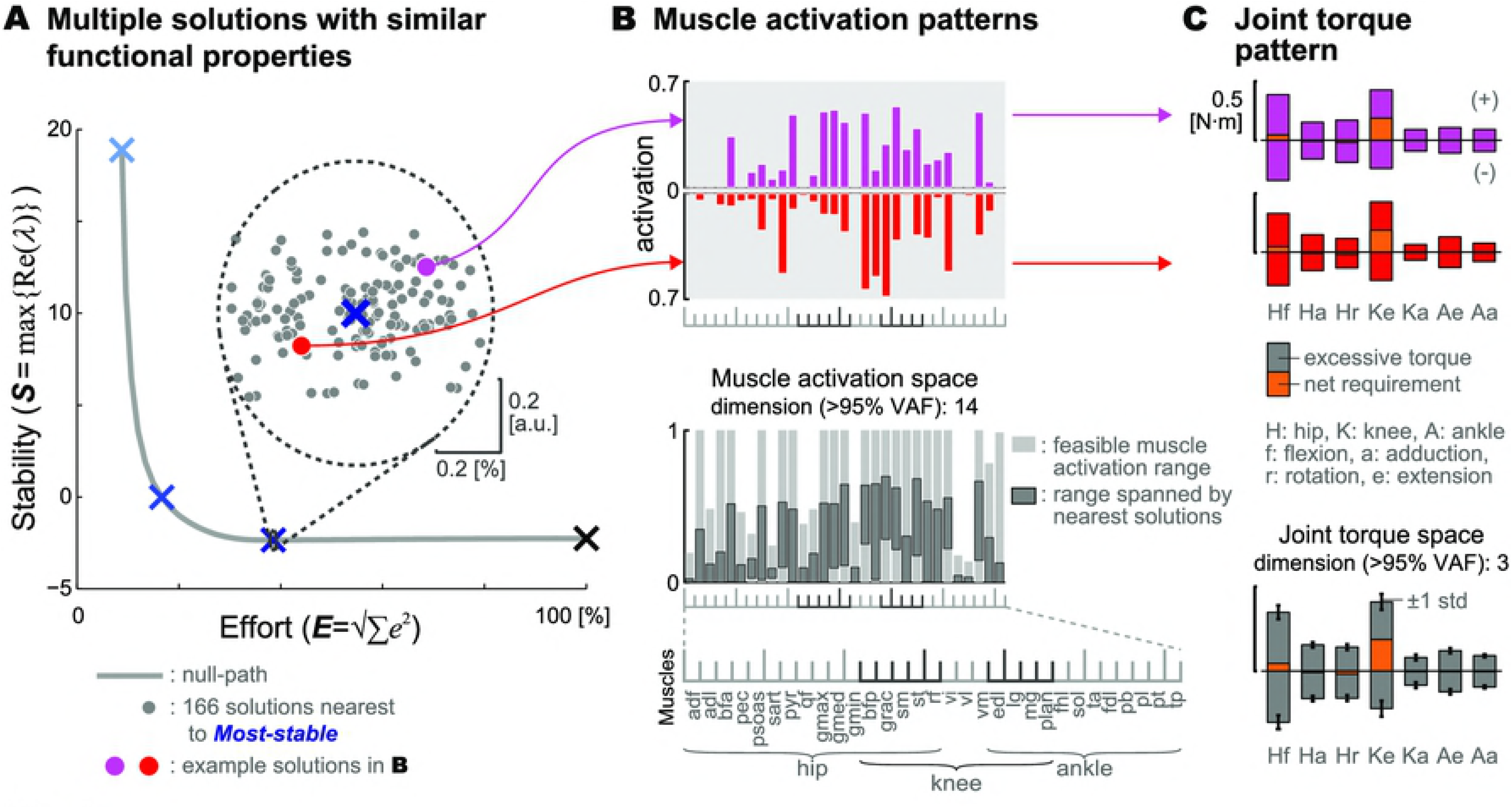
Redundancy among multiple solutions near the most-stable solution. (A) Total 166 solutions (gray dots) nearest to the most-stable solution (‘x’ in dark blue) on the null-path were selected. (B) Two solutions that were very close to each other in functional property space had different muscle activation pattern (top). Overall, activation levels of 166 nearest solutions spanned substantial portion of the feasible muscle activation range (bottom, area filled in dark gray). Dimensionality, defined as the number of principal components required to explain over 95% of variance, was 14, indicating substantial redundancy in muscle activation space. (C) Joint torque pattern about the 7 degrees-of-freedom (DoF) for two examples solutions in (B) are shown on top, mean±std across all 166 solutions shown on bottom. Net required torque about each DoF is shown with area filled in orange. Excessive joint torque produced by both agonistic (+) and antagonistic (-) muscles is shown with area filled in red and magenta for two example solutions in (B), and in gray for all solutions. Excessive joint torque pattern was similar across all 166 solutions, with low dimensionality requiring only 3 principal components to account for >95% variance.

We did not find significant constraints on the dimension of muscle activation patterns producing equivalent function. Overall, activation level of each muscle among these 166 solutions spanned 37.4±13.0% (mean± std) of the feasible range across 31 muscles (Fig 4B bottom, area filled with dark gray). Redundancy in muscle activation space was further demonstrated by dimensionality, where at least 14 principal components were needed to account for over 95% of variance among activation patterns of the 166 solutions. Moreover, 100% of variance could be explained by 24 principal components, indicating that these patterns spanned all possible directions in the null space, which will be reduced otherwise, e.g. if constraint on particular pattern of co-activation were to be imposed [40].

All of the 166 nearest solutions achieving equivalent stability produced the same patterns of joint co-activation torque at all DoFs (Fig 4C). Multiple solutions that differed in muscle activation pattern (e.g. two patterns in Fig 4B, top) still exhibited similar joint torque pattern (two example solutions in Fig 4B on top, mean± std across all solutions on bottom), as quantified by the amount of excessive joint torques produced by both agonistic and antagonistic muscles about each DoF (Fig 4C, area filled with red, magenta, or gray), in addition to the net required joint torque to satisfy the task (Fig 4C, area filled with orange). Compared to relatively high dimensionality in muscle space (14), only 3 principal components were required to explain >95% of variance in excessive joint torque patterns produced by the 166 solutions.

### Global maximum-stability and Pareto front

We performed additional searches to identify the Pareto front which defines the solutions exhibiting an optimal trade-off between effort and stability (Fig 5A). The Pareto front contained the minimum-effort solution (Fig 5A, ‘x’ in light blue) and the global maximum-stability solution (***S***=-3.56 and ***E***=44.8%; Fig 5A, ‘x’ in green), while lying below the null-path. In contrast to solutions on the null-path, solutions on the Pareto front were not linear combinations of each other, indicating that such optimality in effort-stability trade-off may not be accessible by simple linear variations along the null space. Beyond the global maximum-stability solution, convex boundary generated using all solutions identified in this study connected to the maximum-effort solution (Fig 5A, ‘x’ in black) with small positive slope, indicating that immediate increase in stability is not possible near the maximum effort region.

**Fig 5.**
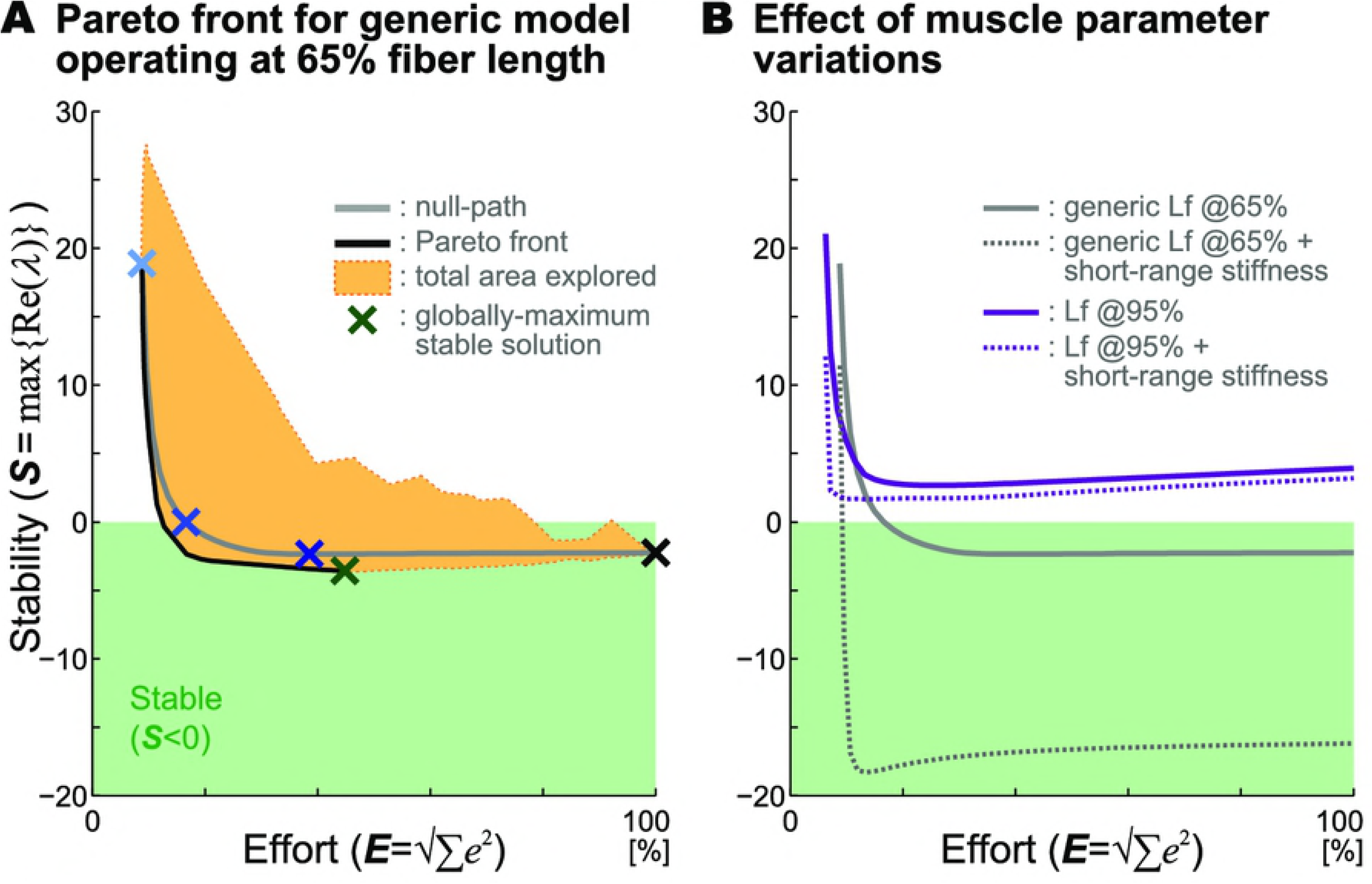
Pareto front of generic model and effect of muscle parameter variations. (A) Pareto front (black solid line) manifesting the optimal trade-off between effort and stability was empirically revealed by first identifying the global maximum-stability solution (‘x’ in dark green) with a heuristic search, and using all solutions explored in this study (area encompassed in orange). The shape of the Pareto front closely followed the null-path (gray), but was consisted of solutions that were more stable than the null-path. (B) Muscle parameters were varied to investigate how other intrinsic stabilizing mechanism and altered condition in force-length relationship affect the mapping in the functional property space. Compared to the generic model with muscles operating at 65% normalized fiber length (solid gray), adding short-range stiffness (dotted gray) drastically improved stability for all solutions on the null-path. Solutions on the null-path for the model with muscles operating at 95% normalized fiber length i.e., plateau region of force-length curve, was always unstable (purple), which could not be stabilized even with short-range stiffness (dotted purple).

### Effects of muscle parameter variations on functional properties

Altering intrinsic muscle properties changes the limb stability but not the qualitative effort-stability trade-off relationship (Fig 5B). Stability drastically improved when intrinsic stabilizing mechanism such as short-range stiffness was added to the generic model with fibers operating at 65% normalized length (Fig 5B, dotted line in gray). On the other hand, the null-path shifted towards unstable region (**S**>0) when muscle fibers operated at plateau region (e.g. 95% normalized fiber length) of the force-length relationship curve (Fig 5B, solid line in purple). The 95% model could not be stabilized, even with addition of short-range stiffness (Fig 5B, dotted line in purple), suggesting that greater force capability offered by operating at near optimal fiber length may come at the cost of minimal stabilizing effect intrinsically provided by force-length relationship.

## Discussion

Our study reveals a wide range of feasible muscle activation patterns for performing a motor task that can vary dramatically in effort and stability, providing insight for interpreting individual variations in experimental muscle activation patterns as well as for generating more physiological simulations of movement in both health and disease. We show that a small increase in muscle coactivation can dramatically increase the mechanical stability of a limb and likely reflect more physiological motor solutions that are necessary for relatively slow sensorimotor feedback pathways to be effective. Further, musculoskeletal simulations using minimum-effort solutions are unstable, and can be easily stabilized by small increases in muscle coactivity within the task nullspace, making simulations more physiological and facilitating computations involved in optimization. We also found that larger amounts of muscle coactivation, such as those found frequently in motor impairments, do not confer additional limb stability, and may in fact impede the ability to improve function through variations in muscle patterns. Our simulations also demonstrate a diversity of functionally-equivalent muscle coordination patterns, which may underlie individual differences in motor solutions with similar behavioral outputs. Overall, our study is the first to systematically characterize the stability and effort trade-offs of functional muscle activation patterns, providing a theoretical basis for individual differences in muscle coordination and how they may affect motor learning and rehabilitation. Further, our methods for searching redundant muscle coordination patterns provide a novel computational framework for generating more appropriate musculoskeletal simulations for understanding both normal and impaired motor control.

This is the first simulation study to explore how muscle co-activation affects both muscular effort and limb stability to explain biological variability in motor patterns. In general, we demonstrate a reciprocal relationship between effort and stability, however, high effort does not automatically confer stability. By mapping out the landscape of effort and stability, our work provides insight into properties of redundant motor solutions within the task nullspace, and the implications for individual variations in motor patterns. Further, this landscape provides a framework for understanding how the nervous system may converge on different redundant motor solutions in learning, disease, and rehabilitation. Our work builds on previous studies demonstrating that increasing muscle co-activation can increase the instantaneous stability of the limb in response to perturbations and changes in limb configuration [36, 40], by explicitly examining the relationship between effort and stability. Further, we took advantage of the finding that maximal effort solutions in arm postural control increases limb stability [39] by comprehensively exploring the continuum of solutions along the null-path between the minimum and maximum effort solutions. While we did not explicitly study effects of limb configuration [56-58], our prior work showed that non-minimum efforts solutions are more likely to generalize their function across different postures, potentially making them more robust for control [59]. Such robust properties of motor solutions could explain our prior observations that individual differences in motor module, a.k.a., muscle synergy, patterns are observed across limb postures and for different tasks [1, 2, 9, 60, 61].

Our results suggest that humans may use low-, but not minimum-effort solutions for motor control due to the need to stabilize local joint dynamics. Increasing co-activation is an intuitive feed-forward mechanism of stabilization in many motor tasks, such as maintaining limb [24, 62] or trunk posture [51, 63], or reaching [64, 65]. Consistent with prior studies [36, 37, 39, 40], we show that minimum effort solutions are very unstable. In our simulation particularly, perturbation doubling time for the minimum-effort solution was on the order of tens of milliseconds, contrary to very small initial kinematic changes observed experimentally during this period [66, 67]. While our minimum effort solution was about 9% maximum effort, an increase to 16% maximum effort dramatically increased the stability of the system such that little change in joint angle would be expected. Further, maximum stability was achieved at about 40% maximum effort. The use of low-levels of muscle co-activation in healthy humans is further supported the observation that long-term training can decrease nominal levels of muscle co-activation, as observed in highly-trained dancers [3]. Rather than being at an absolute optimal solution in terms of effort, the additional criteria of local stability may play an important factor in the selection of motor solutions.

We provide a simple method for identifying more robust and physiologically-relevant motor solutions for musculoskeletal simulations. Here we show that linear combinations of the maximum-and minimum-effort solutions found through static optimization [68] provide a continuum of muscle activation patterns that lie near the Pareto front in terms of maximizing stability for the least amount of effort. More stable solutions within the task nullspace could be easily found by adding a weighted term in a cost function without the need to explicitly compute the Lyapunov stability. While minimum-effort solutions estimated from musculoskeletal models often generate major features of muscle activity, these solutions are fragile and likely to be dynamically unstable in response to perturbations, particularly for motor tasks such as standing and walking [37, 69]. Our results suggest that a small increase in effort could greatly increase the stability of the simulated musculoskeletal system, reducing the sensitivity to small errors due to differentiations methods and discretization [38, 70], improving the stability of forward dynamics simulations. Our prior work demonstrates that the Lyapunov stability conferred by muscle activation patterns based on linearized system dynamics are predictive of the response of musculoskeletal simulations to perturbations during fully nonlinear forward dynamics simulations [36]. Increasing the stability of computed motor solutions in simulations could increase the efficiency of forward dynamics solutions of movement [71, 72] or global optimizations [73, 74]. Further, such musculoskeletal stability is also critical for simulating systems with sensorimotor feedback, which in humans have control delays on the order of 50-150 ms [29] that are often ignored in simulations [75-79].

The stabilizing properties of different muscle activation patterns arises from intrinsic muscle properties such as force-length and force-velocity relationships as well as muscle short-range stiffness. We used Hill-type muscle models operating at 65% optimal length to maximize stability from the muscle force-length relationship. Although solutions did not achieve absolute stability (max real part of eigenvalue <0) when operating near optimal length (e.g. 95%, plateau region), as often observed in many motor tasks in humans in animal [80-82], it did not change the overall result that a small increase in effort drastically improve stability where perturbation doubling times would increase dramatically. For example, at 95% optimal length, solutions with over 10% maximum effort would exhibit relatively small change in posture for ~100ms following a perturbation. Adding muscle short-range stiffness [83] further improve stability to discrete perturbations at both 65% and 95% optimal length, but did not make the model at 95% optimal length explicitly stable in terms of Lyapunov stability. It is possible that a different model of short-range stiffness model [66] could further increase postural stability.

The Lyapunov stability metric may be important for comparing the properties of different muscle activation patterns used in simulations. We studied a quasi-static task in which a single state-transition matrix can be found, the method can be easily extended to movement trajectories. When a muscle activation pattern has been identified (e.g. static optimization or computed muscle control [34]) at a given time point, the linearized state transition matrix will predict the rate (i.e., eigenvalues) and the kinematic mode (i.e., eigenvectors) at which the system states evolve with respect to this equilibrium in presence of small perturbation [84, 85]. Such predictions provide a principled method to evaluate and compare the effects of muscle coactivation patterns on intrinsic stability in any movement, which are not evident in simulated kinematics or external kinetics. While it may be difficult to explicitly include Lyapunov stability as a term in a cost function because the solution space is highly nonlinear and nonconvex, the Lypaunov stability could still be a generalizable tool and metric for understanding the functional role of muscle co-activation in health and disease.

While small increases in muscle co-contraction can be useful for stabilizing musculoskeletal dynamics, high levels of co-contraction–as observed in motor impairment–may come at the cost of excessive effort as well as increased difficulty in improving motor function through rehabilitation. Increased muscle co-activation during movement is observed in people with neural motor impairments [32] and orthopedic impairments including low back pain [47, 48], as well as in older individuals without a clinical diagnosis who experience diminished mobility due to aging [86]. When treating these populations, clinicians often experience difficulty in training individuals with motor impairments to replicate more efficient motor patterns [87-91]. Accordingly, our simulations showed that varying muscle activation patterns around high-effort solutions typically resulted in little change in either effort or stability. As such, the difficulty in finding lower-effort solutions through training and/or rehabilitation may stem in large part due to the inherent properties of redundant high-effort solutions within the task space, which are then further confounded by neuromuscular constraints limiting changes in muscle coordination patterns in neurological disorders such as stroke [46, 92].

In conclusion, the landscape of trade-offs between muscle effort and limb stability could underlie diversity in muscle activation patterns observed across individuals, disease, learning, and rehabilitation. A redundant solution space provides a multitude of muscle activation patterns that have similar functional properties, and individuals may exploit such variability to discover new patterns during motor learning [93-95]. Similar exploratory processes of motor learning have been shown in studies with other species such as song birds [96, 97], rodents [98] and primates [99, 100]. This implies that variability in muscle activation patterns cannot be simply attributed to noise inherent in the sensorimotor system, but may reflect individual differences in selection amongst the abundance of functionally equivalent motor solutions [101]. For example, the “bumpiness” of the landscape may explain why habitual versus optimal patterns are preferred [43-45]. Although we used two functional properties of effort and stability, other multi-objective criteria can be used such as minimizing trajectory error, energetics [102], and ability to generalize [59, 94, 103] or switch [104] across motor tasks. Moreover, the multitude of solutions and local minima may support the observation of individual-specific patterns of muscle co-activation that are regularly observed during gait and balance tasks in healthy humans and animals [1, 2, 9, 60, 61, 105]. Rather than being at an absolute optimal solution in terms of effort, the additional criteria of local stability suggest that individuals use “slop-timal” [106, 107], or good-enough [53] motor solutions.

## Materials and methods

### Musculoskeletal model and target endpoint force

We used a detailed musculoskeletal model of a cat hindlimb [108] to examine muscle activation patterns to produce an experimentally observed endpoint force during postural response in a cat [9] (Fig 1A). Details of this three-dimensional model are described elsewhere [13, 40]. Briefly, the model (Fig 1B) included seven rotational degrees of freedom at the anatomical joints (hip flexion/extension, hip abduction/adduction, hip rotation, knee extension/flexion, knee abduction/adduction, ankle extension/flexion, ankle abduction/adduction) and 31 Hill-type muscles (name and abbreviation in Table 1). The pelvis was fixed to the ground and endpoint of the limb defined at the metatarsophalangeal joint was modeled as a gimbal joint. Model posture was matched to experimentally-measured kinematics of a cat during quiet standing [109]. The equations of motion describing the dynamics of the limb using joint angles as generalized coordinates 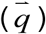 can be given as:

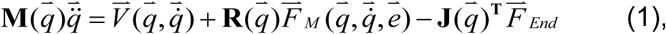

where 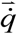 and 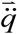 are joint velocity and acceleration vector respectively; 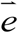 is a vector of muscle activation; **M** is the inertia matrix; **R** is the moment arm matrix; **J** is the endpoint Jacobian; 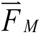 is a vector of muscle forces; 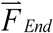 is the target endpoint force vector; 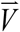 is the Coriolis force vector. Note that the model corresponds to generation of endpoint forces based on no background activity and in the absence of gravity, similar to previous models examining the feasible forces that can be generated by a limb [109-111].

We defined a linear mapping from muscle activation vector to the net joint torque vector required to produce the target endpoint force vector to represent this model at static equilibrium 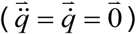:

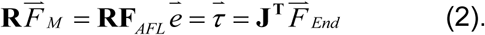

Note that muscle force vector 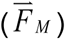 is further factored into a diagonal scaling matrix for isometric force generation (**F**_*AFL*_) based on the force-length relationship [112], multiplied by muscle activation 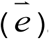. All muscles were set at 65% optimal fiber length in ascending region of the force-length relationship curve [113] to avoid inherent instability owing to lack of intrinsic stiffness in muscles [40].

For the target endpoint force vector, we used an extensor force vector (Fig 1A) that was experimentally measured in active response of a cat following translational support perturbation [9]. This force vector represented the change in the ground reaction force from the background level, averaged over 120-200 ms following the perturbation [30], in which posture of the cat could be approximated quasi-static [67].

### Metrics for effort and stability

In order to evaluate functional properties of redundant muscle patterns that generate the same endpoint force and to map a functional property space in terms of effort and stability, we defined quantitative metrics for each criterion. We defined the metric for effort (***E***) to be the square-root of sum of squared activations (Eq 3), which is equivalent to summing muscle stress [15, 68]:

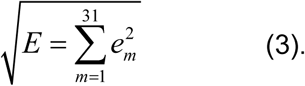

To normalize the level of effort across different muscle activation patterns, we identified the global minimum-effort solution 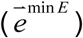 and maximum-effort solution 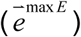 for the static mapping (Eq 2), using quadratic programming. The effort for any muscle activation pattern examined in this study was then normalized to percent of the global maximum, i.e., ***E*** of the maximum-effort solution.

We defined the metric for stability using Lyapunov stability of the linearized model. The full nonlinear system (Eq 1) was linearized about a static equilibrium point, defined by a muscle activation pattern that satisfies the endpoint force generation (Eq 2), using software *Neuromechanic* [114]. Specifically, the system equation incorporated joint torques generated by muscles:

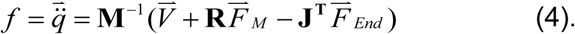

The system was linearized by numerically computing the partial derivatives with respect to kinematic states using first-order Taylor-series expansion to obtain the state space representation:

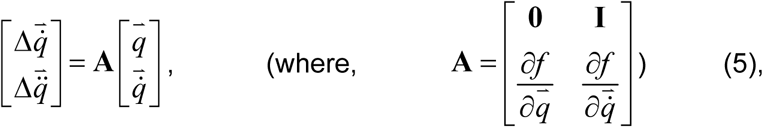

The state matrix (**A**) was used to calculate the eigenvalues (*λ*) of the linearized system. For a given muscle activation pattern, the metric for stability (***S***) was defined as the maximum real part of the 14 eigenvalues of **A** such that

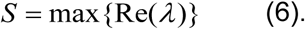

We only considered 8 out of 14 eigenvalues that had the largest real parts in magnitude, which correspond to the 8 modes (or eigenvectors) that are relevant to the dynamics of the system in physiological timescale. Such estimate can be made when converting eigenvalue to a time constant, defined as the doubling time [40]. Due to the constraint that eliminates endpoint translation in 3 directions, effective degrees of freedom of the model is reduced from 7 to 4. As a result, 6 of the 14 eigenvectors are modes that do not affect the dynamics of the system, corresponding to the 6 eigenvalues that are near zero (similar to rigid-body modes) which are very small in magnitude. These 6 eigenvalues typically were less than 1e-5 in magnitude, and thus were not relevant in physiological time scales: the time at which the magnitude of a perturbed response is reduced to 50% is longer than 6.9e4 seconds. In contrast, the 8 eigenvalues that were considered in ***S*** for a given solution were typically larger than 1e-3 in magnitude.

A solution of the system defined by a given muscle activation pattern was determined “stable” if ***S***<0, and “unstable” if ***S***>0. Further, because the magnitude of ***S*** predicts the rate at which a perturbed system will return to (if ***S***<0), or deviate from (if ***S***>0) the equilibrium, a solution is to be “less stable” for greater value of ***S***, and “more stable” for smaller value of ***S***. This metric from system theory, i.e., Lyapunov indirect or linearization method, has been shown to predict the behavior of perturbed nonlinear systems in simulations [36, 40].

### Null-path between the minimum- and maximum-effort solutions

In order to explicitly examine how stability changes along a given direction in the null space across all possible effort levels, we evaluated effort (***E***) and stability (***S***) of 49 intermediate solutions that were evenly spaced between the minimum- and maximum-effort solutions in muscle activation space (Fig 2B). These solutions 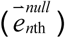 were computed by linearly scaling the difference between the minimum-effort solution 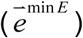 and the maximum-effort solution 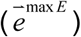:

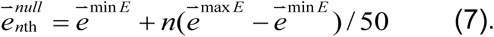

Note that difference between any two solutions that satisfy the torque requirement (Eq 2) belongs to the 24-dimensional null space defined by the linear mapping matrix (**RF**_*AFL*_), i.e., the vector difference will produce zero torque (e.g. 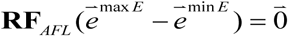. Therefore, solutions generated as above (Eq 7) lie along the “null direction” defined by the vector difference 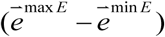. Here, we defined the unique path between the two solutions in the muscle activation space as the null-path.

We mapped the null-path onto the functional property space by evaluating the effort (***E***) and stability (***S***) of each intermediate solution, revealing the projection from the linear muscle activation space to the functional property space. In particular, we were interested in whether the minimum-effort solution, representing the least amount of co-activation for a given task, is unstable, and whether the maximum-effort solution, with highest level of co-activation, is stable. Further, this null-path was used to demonstrate whether an unstable solution can be made stable by following a defined direction in muscle activation space, and to quantify the amount of effort necessary to make the minimum-effort solution stable.

To quantify the sensitivity of stability to change in effort, we calculated instantaneous slope of the null-path at each of the 51 solutions by numerically differentiating the cubic curve generated by interpolating the 51 points on the null-path using small intervals (0.01%) in terms of effort.

### Exploring the neighboring solutions

In order to reveal the local landscape of the solution space with respect to effort and stability, we explored the *neighboring solutions* around the null-path 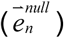 between the minimum- and maximum-effort solution, and mapped them onto the functional property space. The *seed* 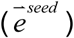 for each exploration was defined as each of the 51 solutions on the null-path (Fig 2A): the minimum- and maximum-effort solutions, as well as 49 intermediate solutions). We defined *neighboring solutions* 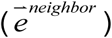 as the solutions that were pseudo-randomly distributed around the *seed* in muscle activation space in which the amount of change, either positive or negative, in any muscle from the *seed* was constrained to be within a given *step size* specified for each muscle:

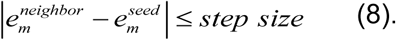

The *step size* was defined for each muscle as a percentage of the feasible muscle activation range (Sohn et al., 2013) for the target endpoint force production, so that the amount of change in any muscle is normalized. To vary the extent to which area in the functional property space is explored by the neighboring solutions around a given seed solution, we used step sizes of 5, 10, and 25% of the feasible muscle activation range.

We generated 262 neighboring solutions 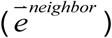 for each seed and step size using the following four steps. In summary, our goal was to explore the functional property space in every possible direction around a given seed, reaching the maximum extent in distance for a given amount of changes allowed in the muscle activation space. To this end, we generated sets of solutions that were randomly distributed around a given seed with varying distance in muscle activation space controlled by the step size:

1. In order to induce random deviations to muscle activations near each of the seed solutions, we first generated 200 perturbed patterns 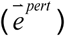, in which each element in 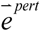, i.e., 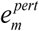, was randomly drawn from a normal distribution with mean (*µ*) at activation level of the seed 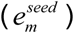 and variance (*σ*^2^) equal to the muscle-specific step size (Fig 3B):

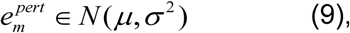

where 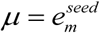 and (σ^2^ = step size. However, in order to examine perturbed patterns in which 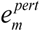 was always within physiological limits (0,1), we limited the range from which the values were drawn using an algorithm for simulating a truncated normal distribution on a finite interval [115, 116], implemented in MATLAB (Mathworks, Inc., Natick, MA, USA). The interval of each muscle was defined as the smaller value of physiological limits (0,1) or two times the step size away from the seed:

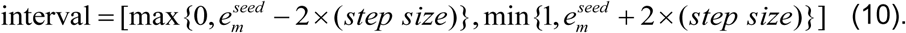
2. We then identified the nearest solutions 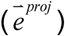 to all 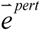 that produced the specified force. To find projections of 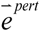 to the solution manifold in a least-squares sense (Fig 3B), we performed optimizations to find muscle activation patterns 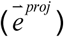 that minimized sum-squared difference to each of the perturbed patterns:

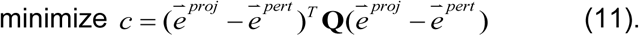

Note that we used a scaling matrix **Q**, which is a diagonal matrix that penalizes the difference between perturbed and projected muscle activation. Elements of **Q** were weighted inversely to the feasible range to prevent projection only occurring in muscles with small feasible ranges. Each optimization was subject to an equality constraint for satisfying the torque requirement:

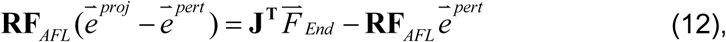

and inequality constraints specifying the search limits 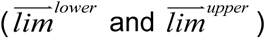 defined either by the step size or the bounds from the feasible range identified for given task:

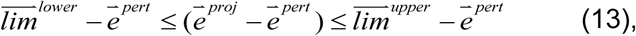

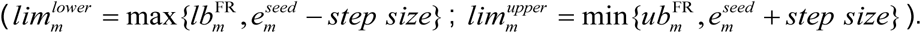
3. To search the functional property space, we defined a new null-path between 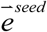 and each 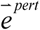, and found extrapolated solutions 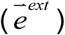 along those paths until muscle activation met the search limits. Null-paths were computed in a same way described above, where the change 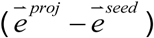 was further extrapolated along the null direction until any of the muscles reached either the lower or upper bound (Eq 13) for given step size. Extrapolated solutions 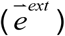 generated this way are true solutions that satisfy the torque requirement, with at least one of the muscle activations lying on the search limits. These were defined as the *neighboring solutions* of the initial seed solution for a given step size. To visualize the path in the functional property space to get to the corresponding neighboring solution, we also computed four intermediate solutions linearly interpolated along the null-path from the seed to each of the neighboring solutions.
4. Finally, in order to encourage the generated neighboring solutions to span the full possible range in muscle activation space for a given step size, we further generated solutions 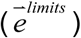 that lie on the search limits (Eq 13). Limit solutions were computed by specifying activation level of each muscle at its lower 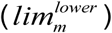 and upper limit 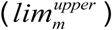, and pushing the other 30 muscles as close as possible to their lower and upper limits. This optimization is similar to that for projected solution (Eq 11-13), where the problem is reduced to solving for a 30-dimensional muscle activation vector that minimizes the distance to either lower or upper limit vector (Eq 13), specified for a given step size. Although solutions generated this way were not guaranteed to satisfy the torque requirement, especially when small step sizes were allowed, more than 57 (60±1.1 across seed and step size) limit solutions were generated for each seed and added to the set of neighboring solutions of the seed for a given step size.

### Characterization of local landscape

We characterized the local landscape around each seed solution using the neighboring solutions mapped in the functional property space. We were particularly interested in whether characteristics of the original seed solution on the null-path are preserved as step size increased. To this end, we identified a 95% confidence ellipse in the two dimensional space of effort and stability (Fig 3C), using the covariance matrix of effort (***E***) and stability (***S***) values evaluated for each set of neighboring solutions, i.e., for each seed and each step size. The eigenvector of the covariance matrix was used to define the major axis of the ellipse, corresponding to the direction in which effort and stability varied maximally, and the minor axis that is orthogonal to the major axis. Note that such procedure is equivalent to performing a principal component analysis. The center of the ellipse was defined at the mean effort and stability values for given set of neighboring solutions. The radii of major and minor axes were defined by scaling the maximum and minimum eigenvalues of the covariance matrix, respectively, with chis-squared value for 95% percentile. Note that while such ellipse may not encompass or be fully occupied by the actual neighboring solutions distributed in the functional property space, our analyses (see below) mainly used the values derived from the covariance matrix and does not depend on geometrical distribution.

We quantified four metrics to examine the extent to which local landscape explored by the neighboring solutions differed across step size. First, to quantify how sensitivity of stability to effort change, we computed the amount of deviation in orientation of the major axes from the slope of the null-path at given seed solution. To avoid numerical error resulting from directly using the values for the slopes (e.g. near zero or near infinity), we converted the difference in angles (degrees), i.e., arctangent of slope, in the two-dimensional functional property space. Second, to quantify the amount of space explored by the local search, we calculated the area of the ellipse, however, simply defined as the multiplication of radii of major and minor axes while omitting the universal scaling constant *pi*. Third, to quantify how far majority of search reach in the functional property space, we computed the Euclidean distance between the seed solution and the center of the ellipse in the functional property space. Finally, to quantify the extent to which local search resulted in exploring solutions that lie orthogonal to the null path, we computed eccentricity of each ellipse:

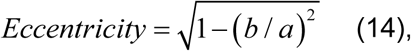

where *a* and *b* are radii of the major and minor axes of the 95% ellipse, respectively. We tested whether above four metrics change as step size increase using a nonparametric Kruskal-Wallis test, where statistical significance was evaluated at α=0.05 with Tukey’s honestly corrections for multiple comparison.

To visualize the distribution of neighboring solutions both in muscle space and functional property space (Fig 3B and 3C, respectively), we used Kernel smoothing function estimate (‘ksdensity.m’ in MATLAB), which was normalized to have maximum value of 1. Note that this estimate was used for visualization purpose only.

### Investigating redundancy within equivalent functional properties

In order to examine the extent to which solutions can vary and still have similar functional properties, we compared multiple solutions that are within a small range of effort and stability (Fig 4). Specifically, we were interested in whether near-maximal stability could be achieved using vastly different muscle patterns. Thus, among all neighboring solutions investigated, we examined solutions that were nearest to the most-stable solution (Fig 4A, ‘x’ in dark blue) on the null-path between the minimum- and maximum-effort solutions. In order to avoid selecting solutions that are inherently similar in muscle activation pattern, we first sorted 15 solutions that were nearest in Euclidean distance to the local maximum stability solution from each set of neighboring solutions around a given seed, across step sizes. Among these 765 solutions (15 from each of 51 seeds), we selected solutions that had effort and stability within 10% and 1% difference, respectively, from the most-stable solution on the null-path. This resulted in total 166 solutions near the most stable solution on the null path (Fig 4A, gray dots).

To quantify the spatial amount of redundancy, we computed percentage of range spanned by the above nearest solutions for each muscle with respect to corresponding feasible muscle activation range. To examine the redundancy in the muscle activation space, we also computed the dimensionality within the activation set [40]. Dimensionality was computed using principal component analysis, defined as the number of principle components required to explain over 95% variance in data (i.e., a 31×166 matrix).

To determine whether similar level of stability was achieved by different types of stabilizing strategy, e.g. via joint stiffening, we examined pattern of co-contraction about the joints. For each of the 166 selected solutions, we computed the amount of excessive joint torques produced by agonistic and antagonistic muscles about each of the 7 DoFs, which were further concatenated in to a single vector:

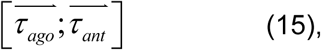

comprised of 7 dimensional column vector for agonistic torques 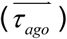 and another 7 dimensional column vector for antagonistic torques 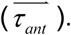 Dimensionality within these 166 patterns of excessive torques was determined as in muscle activation space, defined as the number of principle components required to explain over 95% variance in data (i.e., a 14×166 matrix).

### Global maximum-stability and Pareto front

We found a global maximum-stability and defined the edge that connected the minimum-effort solution and global maximum-stability solution as the Pareto front and compared the values and shape to the null-path between the minimum- and maximum-effort solutions. To examine an explicit trade-off for minimizing effort while maximizing stability, we identified Pareto front that quantifies the maximum level of stability that can be achieved for a given amount of effort, or vice versa, the minimum amount of effort required to achieve certain level of stability. To define this edge, we first performed a heuristic search that identified the global maximum-stability solution (most stable with smallest ***S***). Across all solutions we found the solution with smallest values of ***S*** 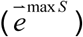 to explore neighboring solutions. If solutions that were more stable was found, it was assigned as the new global maximum-stability solution and the search was repeated with the new seed. Even when no solution more stable than 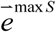 was found in a given iteration, we continued the search using the same until there were no change in maximum stability for 20 consecutive iterations. We then defined Pareto front by identifying the convex outer-most edge (‘convhull.m’ in MATLAB) connecting the minimum-effort solution and the global maximum-stability solution using all of the solutions explored in this study.

### Effects of muscle parameter variations on functional properties

We further investigated how altered condition in force-length relationship and other possible intrinsic mechanisms that contribute to stability affect the mapping in the functional property space. First, stability of the original 51 solutions on the null-path were evaluated in a model implemented with a short-range stiffness in muscles [83, 117] to represent the instantaneous force-length behavior. However, the short-range stiffness model was modified because we assumed tendons to be inelastic in the static mapping (Eq 2). Hence, our short-range stiffness model represented the instantaneous stiffness only from the active muscle, assuming an infinite stiffness in the tendons that were in series. In result, we used a force-relationship curve with a constant slope [83]. In addition, we did not neglect the force-velocity relationship that provided damping as in the generic model because its contribution may be large in the linearization process where we used numerical perturbation method.

Secondly, we computed another set of 51 solutions that constituted a null-path but with a modified model where fiber lengths of all muscles were set at 95% optimal fiber length, i.e., plateau region of the force-length relationship curve [112] and thus more prone to destabilization. Note that these solutions are not the same as the original solutions, because the scaling matrix for isometric force generation (**F**_*AFL*_) in Eq 2 is different. However, each of the 51 solutions in the two models (65% and 95%) were very similar (cosine of the angle between the two muscle activation vectors was 0.998±0.002 across 51 solution pairs). In addition, we also evaluated the null path of the 95% model with added short-range stiffness properties of active muscles using the same method described above.

